# Co-incubation of dsRNA reduces proportion of viable spores of *Ascosphaera apis*, a honey bee fungal pathogen

**DOI:** 10.1101/852699

**Authors:** James P. Tauber, Ralf Einspanier, Jay D. Evans, Dino P. McMahon

## Abstract

There are viral, fungal, bacterial and trypanosomal pathogens that negatively impact the individual and superorganismal health of the western honey bee. One fungal pathogen, *Ascosphaera apis*, affects larvae and causes the disease chalkbrood. A previous genome analysis of *As. apis* revealed that its genome encodes for RNA interference genes, similar to other fungi and eukaryotes. Here, we examined whether *As. apis*-targeting double-stranded RNA species could disrupt the germination of *As. apis*. We observed that when spores were co-incubated with *As. apis*-targeting dsRNA, fewer spores were activated for germination, suggesting an uptake of exogenous genetic material at the very onset of germination and consequent damage to essential transcripts needed for germination. Overall, these results indicate that the causative agent of chalkbrood disease, *As. apis*, can be successfully targeted using an RNAi-based strategy.

## Introduction

Ribonucleic acid interference (RNAi) is an evolved viral defense strategy in eukaryotic organisms which is likened to restriction enzymes in prokaryotes. It is also involved in host gene regulation. As a viral defense strategy, RNAi is a gene silencing mechanism that cleaves viral sequences, thus eliminating translation of the virus in the host. The host’s canonical RNAi-related proteins (“machinery”) include Dicer, Argonaute and RNA-dependent RNA polymerase proteins, and are found in both fungi and animals (Dang, Yang, Xue, & Liu, 2011; Hu, Stenlid, Elfstrand, & Olson, 2013). Double-stranded RNA (dsRNA), a regular component of replication in many viruses, is recognized by Dicer, which cleaves this dsRNA into 21 to 25 nucleotide dsRNA fragments (“diced RNA”). A single strand from these dsRNA fragments guides Argonaute to the respective complementary viral sequence and cleaves this region. RNA-dependent RNA polymerase, when present, can amplify these RNA strands, thereby amplifying the RNAi response.

RNAi studies in insects, in particular the honey bee (*Apis mellifera*), have shown that the RNAi system can successfully be exploited to reduce pathogen titers. The honey bee possesses the necessary RNAi machinery. In the laboratory to activate and manipulate the RNAi response in bees, injection of dsRNA is one route, although feeding bees dsRNA appears to be a simpler and a successful method as well (Yang et al., 2018). To demonstrate the importance of the RNAi antiviral defense response in bees, when honey bee transcripts of *Dicer* were knocked down using dsRNA, researchers observed an increase in viral titers in the host (Brutscher, Daughenbaugh, & Flenniken, 2017). It is important to understand how to combat viral pathogens like Deformed wing virus (DWV) and Israeli acute paralysis virus (IAPV) because such viruses are linked to overwinter colony loss (Döke, Frazier, & Grozinger, 2015). Double-stranded RNA fed to honey bees with complementing sequences to DWV or IAPV led to reduced viral titers in the honey bee (Maori et al., 2009). However, ambiguous dsRNA sequences, including even the non-complementary sequence of *GFP*, can also induce an RNAi antiviral defense or systemic suppression of transcripts in the honey bee (Flenniken & Andino, 2013). RNAi has additionally been used against the parasite *Nosema ceranae*, an emerging honey bee spore-forming fungal parasite that is a significant cause of concern. In one case, dsRNA targeting the polar tube of *N. ceranae*, which is used by *N. ceranae* to penetrate the host’s epithelial cells to initiate its reproductive cycle, led to significantly fewer spores in the honey bee gut (Rodríguez-García et al., 2018).

Interestingly, a host’s own RNAi mechanism can be used against itself. For example, injecting beetles with beetle-targeting dsRNA helped researchers create a catalogue of essential/lethal genes as well as constituting experimental evidence that a gene was connected to a phenotype throughout the insect’s development from egg to adult (Jasrapuria, Specht, Kramer, Beeman, & Muthukrishnan, 2012; Schmitt-Engel et al., 2015). As noted above, fungi possess the necessary components for RNAi, in addition to other honey bee-associated pathogens. The opportunistic/pathogenic, spore-forming, ascomycetous fungi, *Aspergillus* and *Fusarium*, are perhaps the best studied organisms where researchers used RNA species to activate the RNAi mechanism in the fungus and thereby against itself (Baldwin et al., 2018; Kalleda, Naorem, & Manchikatla, 2013). In this system, for example, germinating *Aspergillus nidulans* was introduced to diced RNA species that targeted putative Ras family genes for which some of the respective proteins are completely essential for growth and development (Kalleda et al., 2013). A knockdown (and not knockout) of Ras transcripts led to reduced fungal growth. In this study, no special transfection method was done to transfer RNA into the cells, meaning that fungi can, in principle, actively uptake exogenous genetic material. This was in alignment with other studies that showed growing fungi take up exogenous genetic material and transfection reagents are unnecessary. However, increased effectiveness of gene silencing methods from the use of such reagents may be areas for future improvements regarding this technology. In *A. fumigatus*, the uptake of exogenous RNA was recorded over a period of two hours during germination (Jöchl, Loh, Ploner, Haas, & Hüttenhofer, 2009). In another model organism, *Pecoramyces ruminantium* (Division: Neocallimastigomycota), the uptake of exogenous dsRNA by germinating spores occurred within just 15 minutes after its introduction to the activated spores (Calkins et al., 2018).

*Ascosphaera apis* (*As. apis*) is a fungal pathogen of the honey bee that causes the disease chalkbrood (reviewed in (Aronstein & Murray, 2010)). Spores of this fungus are ingested by larvae which then germinate in the gut where the anaerobic environment activates/stimulates spore germination. When the spore is activated for germination, it undergoes physical changes such as swelling and then the emergence of a primary germ tube. During outgrowth in the larva, the hyphae breaches the gut epithelial cell wall, followed by the larval body, altogether leading to the visual characteristic of a “mummified” larva. This does not occur in adults. The *As. apis* genome is known to encode the RNAi machinery (Cornman et al., 2012). While chalkbrood may currently be less concerning than other emerging infectious honey bee diseases such DWV or *Nosema*, *As. apis* can nonetheless reduce brood levels in infected colonies. Particularly in colonies already weakened by infections with other pathogens, chalkbrood can potentially and additively threaten the long-term survival of these colonies. This is in addition to other possible effects from other filamentous fungi (*e.g*., Stonebrood disease; (Jensen et al., 2013) or yeasts (Tauber, Nguyen, Lopez, & Evans, 2019)).

As honey bees are affected by a multitude of stressors and pathogens, it is important to seek remedies against all pathogens. To this end, we hypothesized that RNAi may be applicable against *As. apis* as described in other fungi. Activated *As. apis* spores were introduced to Ras family and also cell wall synthesis targeting dsRNA which we presumed to be essential for growth and development. We observed a reduced number of germinating spores when *As. apis* was exposed to these *As. apis*-targeting dsRNA. We believe that this establishes a platform for exploiting the RNAi machinery of *As. apis*, and possibly synergistically with that of the honey bee, to combat this pathogen as well as to discover, beyond sequencing projects, virulence factors that this fungus utilizes during host infection.

## Materials and Methods

### Fungal strains, growth and spore production

Two mating-compatible, species-confirmed *Ascosphaera apis* strains (AFSEF 7405 and ARSEF 7406) were ordered from the ARS Collection of Entomopathogenic Fungal Cultures, USDA-ARS (NY, USA) as actively growing cultures. An agar plug containing *As. apis* mycelia was plated onto potato dextrose 1.8% agar (Roth) to make working plates, and the plates were sealed with PARAFILM^®^ M, wrapped in aluminum foil and grown at 30°C under aerobic conditions. One agar plug from each mating strain working plate that contained actively growing cultures was then placed across from one another on a similar plate, which was incubated, as described in (Jensen et al., 2013), in order to produce spores. After three to four weeks of growth, the interaction zone between the two strains appeared pigmented, indicating the production of spores. The spores generally do not activate or germinate on agar after they are produced without CO_2_ stimulation. About 1 ml of autoclaved dH_2_0-tween (one droplet of Tween^®^ 80 per 500 ml water) was added on top of the interaction zone, the spores scraped into the solution and then the solution was collected into a sterile plastic centrifuge tube. This was repeated on the same plate. Multiple plates were also incubated in parallel. The spore suspensions were grounded using a sterile plastic pestle. This suspension was filtered through a sterile folded cheese cloth. The spores were mixed with 50% sterile glycerol in dH_2_0 to create a 25% glycerol stock and the concentration of spores were then counted in a Neubauer improved chamber (BLAUBRAND^®^) using 10 μl of the stock. The stock was stored at -70°C until thawed as needed for experiments.

### Cloning and dsRNA production

Our original goals were to i) modify plasmid L4440 and produce dsRNA in *E. coli* (Solis, Santi-Rocca, Perdomo, Weber, & Guillén, 2009); ii) produce dsRNA that targeted putative Ras family genes and other genes like cell wall synthesis that are likely essential/lethal, as done in *Aspergillus nidulans* (Kalleda et al., 2013); iii) use the RNAi machinery of the fungus rather than the bee host to induce an RNAi effect, similar to *A. nidulans* and other filamentous fungi (Cornman et al., 2012; Dang et al., 2011); and iv) introduce dsRNA to the spores as-is because we presumed that germinating spores will uptake or bind to exogenous genetic material upon CO_2_ activation and without the need for transfection reagents, as observed in other fungi and microorganisms (Calkins et al., 2018; de Sousa Pereira, Piot, Smagghe, & Meeus, 2019; Jöchl et al., 2009). To this end we searched the most recent *As. apis* ARSEF reference genome (Shang et al., 2016) for predicted Ras family and glucan synthesis proteins using keywords within NCBI. We selected the following proteins: ‘Ras GTPase-activating-like protein rng2’ (KZZ95061.1; Ras gene) and ‘1,3-beta-glucan biosynthesis protein’ (KZZ95811.1; beta-glucan gene). When aligning the primary protein sequences of these two proteins in NCBI’s blastp there was 35% coverage with 46.81% identity. To clone partial sequences of these genes, total RNA was isolated from *As. apis* mycelia from an actively growing culture using the Quick-RNA™ Miniprep Kit (Zymo Research). Complementary DNA (cDNA) was produced using ProtoScript^®^ II Reverse Transcriptase (NEB) for 2 hours at 48°C from an aliquot of the total RNA isolate and priming with d(T)_20_VN; otherwise the manufacturer’s recommend reaction mixture was used. The cDNA was diluted 1:1 with nuclease-free water and used in Q5^®^ High-Fidelity DNA Polymerase (NEB) reactions (as recommended by the manufacturer) using the following primer pairs (listed in the 5’ to 3’ direction). Primers were developed within the online tool Primer-BLAST (Ye et al., 2012) using the database’s cds sequence from the two genes (Ras gene: F: CCTGCAATGGTTTGTGGACG, R: ATTCAGCCTGAGGTTCCTGC, expected: 422 bp; and beta-glucan: F: TTGGCGTGGGTACTTCTTCC, R: TTTACCGTTGTTTGCGCCAG, expected: 542 bp). The partial gene was then checked against the current honey bee genome (GCA_003254395.2 Amel_HAv3.1) within NCBI. An aliquot of the PCR mixture was run in a 1.0% agarose gel (Agarose NEEO Ultra-Quality (Roth)) in 0.5 × TAE buffer to check that the amplicons were of proper size by molecular weight. Our amplified cds sequence of the beta-glucan gene showed one misannotated intron after sequencing (described later), but initially noted by a different expected bp size in the gel (actual amplified length = 455 bp). The remaining PCR mixture was purified using the Monarch^®^ PCR & DNA Cleanup Kit (NEB) and the DNA eluted with nuclease-free water. An aliquot of the purified amplicons was phosphorylated using T4 Polynucleotide Kinase (NEB) using the manufacturer’s recommended reaction mixture and we extended the incubation. In parallel, the plasmid L4440, which was propagated in *E. coli* Mach1™ (Thermo), was isolated from *E. coli* using the QIAprep^®^ Spin Miniprep Kit (Qiagen) and the DNA was eluted with nuclease-free water. The plasmid was cut and de-phosphorylated simultaneously using 1 U Shrimp Alkaline Phosphatase (NEB) and 20 U EcoRV-HF (NEB) in the same reaction mixture. The dephosphorylated/restriction enzyme modified plasmid and phosphorylated amplicons were mixed and the ligation reaction was carried out using T4 DNA ligase (NEB) as instructed on NEB’s website. After overnight ligation, the ligation mixture was incubated with chemically competent *E. coli* HT115 cells (prepared with 100 mM CaCl_2_, 1% PEG 6000, 15% glycerol and dH_2_0), transformed by heat shock and the cells plated on LB-1.8% agar-ampicillin-tetracycline plates (incubated 37°C). Single colonies were selected the next day, inoculated in similarly amended liquid LB and incubated with shaking. Transformation efficiency was very low and resulting in only a couple colonies per transformation; however, almost every transformant contained the plasmid with the insert. The next day, the plasmid was isolated from the dense culture using the Monarch^®^ Plasmid Miniprep Kit (NEB) and the DNA was eluted with nuclease-free water. The insert was checked by Sanger single primer sequencing at Eurofins Deutschland (Germany) using published primers “M13/pUC Forward” (CCCAGTCACGACGTTGTAAAACG) and “L4440 Reverse” (AGCGAGTCAGTGAGCGAG) (Addgene plasmid # 1654; http://n2t.net/addgene:1654; RRID:Addgene_1654). Sequences were analyzed in Serial Cloner (v2.6; Mac). All primers were ordered desalted from Metabion (Germany).

After several attempts to produce large quantities of dsRNA in transformed *E. coli* HT115 and to remove endogenous RNA species from the suspension, we then chose to produce the dsRNA *in vitro*. To this end, we used the aforementioned primers M13/pUC Forward and L4440 Reverse and the isolated plasmid (template) in a OneTaq^®^ 2X Master Mix with Standard Buffer (NEB) reaction as recommended by the manufacturer. The predicted amplicon contained the gene of interest, the flanking T7 promoter sequences, interspersed plasmid nucleotides and the primer sequences (adding 191 bp to the *As. apis*-target sequence). The amplicons were incubated in a 50 μl T7 RNA polymerase reaction at 37°C overnight. The reaction consisted of 1 × RNAPol Reaction Buffer, 1 mM of each rNTP (NEB), 1.25 U RnaseOUT™ (Thermo), 5 mM DTT (Thermo) and 250 U T7 RNA polymerase (NEB). The next day, 6 μl 10 × Dnase buffer, 2 μl amplification-grade DNase I (Invitrogen) and 2 μl nuclease-free water were added to the T7 reaction mixture, and the mixture further incubated for 45 minutes at 37°C to remove DNA. We then used SureClean Plus (Bioline) to purify the dsRNA as this purification method is a “non-toxic alternative” and probably does not have any residual toxic chemicals (*e.g*., as potentially with TRIzol^®^), an important consideration when using the suspension for experiments with an organism (personal communication with the company). We followed the recommended SureClean Plus protocol except that the centrifugation steps were done at 15,000 rpm and for 15-30 minutes, we used 75% ethanol, we did not vortex (only flicking) the RNA and the large white pellet was suspended in 50 μl sterile dH_2_0. An aliquot of the T7 transcription reaction (before and after purification) was run in a 2.0% agarose gel using GelRed^®^ Nucleic Acid Gel Stain (Biotium) as above to confirm the production of dsRNA. We used a 100 bp dsDNA (not dsRNA) ladder (Fermentas); therefore, the size in bp could only be estimated. The concentration of RNA was measured by Nanodrop (Thermo). The purity had acceptable values (A_260/280_ > 2.0 and A_260/230_ > 2.0 in all cases after purification). The dsRNA was used fresh for the experiment as well as from an one-time frozen-thawed aliquot. A mock water SureClean Plus extraction was used for control treatments and this extraction did not produce any bands in the gel nor a spectral readout from the Nanodrop.

### As. apis in vitro tests

To start the *in vitro* tests we first thawed one aliquot of the glycerol spore suspension. To uniformly distribute an equal number of spores per sample, we first made a master mix of the spores, medium and antibiotic (as the preparation was done outside of a laminar flow hood), and then vortexed the mixture frequently while aliquoting the 49 μl spore master mix per 700 μl tube, after which the tubes were randomly assigned a sample code. The final reaction mixture contained 20 μl of the *As. apis* glycerol spore stock, 24 μl of Sabouraud Dextrose broth (Roth), 5 μl of ampicillin-salt (50 μg/mL final concentration, Carl Roth) and approximately 30-32 nM dsRNA (that is, approximately 0.7 to 1 μl of the dsRNA-water solution amended with sterile water to 1 μl when required). A water suspension from a mock SureClean Plus extraction was used *in lieu* of dsRNA for the positive and negative controls. In total, there were five treatments: a positive control (mock extraction and + CO_2_), a negative control (mock extraction and - CO_2_), *As. apis* with Ras gene dsRNA (+ CO_2_), *As. apis* with beta-glucan gene dsRNA (+ CO_2_) and an unrelated dsRNA control (+ CO_2_), and the treatments were always run in parallel with one another per trial and in duplicate or quartet. For the unrelated dsRNA control, we used a non-ribosomal peptide synthetase-like gene (*NPS3*) from an unrelated fungus, the basidiomycete *Serpula lacrymans* (a reference strain at the BAM Institute), and this gene shares no similarity with genes from *As. apis* by blastn. The molecular techniques to create the *NPS3* dsRNA control were exactly like the procedures for the other partial genes (EGO23141.1: F: TTCCGTGCTCGTGGATTTGA, R: AGCTTCTCCGGTGTCAACTG, expected: 419 bp).

It is well established that a flush of CO_2_ activates spores and it is the most reliable way to active spores for germination (Heath & Gaze, 1987). Therefore, we placed the sample-coded tubes with their lids opened and in a randomly assorted array into a sealable plastic container and flushed the container for 15 seconds with CO_2_ (Linde Gas). The tubes with opened lids were incubated in the container for 15 minutes, a modified version of the ‘flush method’ set out in (Heath & Gaze, 1987). This was followed by one additional CO_2_ flush and then the tube lids were immediately closed. The tubes were placed at 30°C (at this point, CO_2_ concentration was not measured and the spores were exposed to light). A remaining aliquot of prepared spores was examined by microscopy at the onset of the experiment; in each trial we observed that the spores at the beginning of the experiment were unactivated and appeared like the negative control throughout the experiment (see Results).

Approximately 24-26 hours later, spores were counted as carried out before for *Ascosphaera apis* (Heath & Gaze, 1987) and *Ascosphaera aggregata* (Goettel, Duke, Schaalje, & Richards, 1992) germination counts. The tube was vortexed for 5 seconds, the solution pipetted up and down three times and then 10 μl was loaded into a Neubauer improved chamber and allowed to settle for two minutes. Counting of activated and unactivated spores was observed at 400 × with phase contrast. Unactivated versus activated spores were easily distinguished from one another using two observations: 1) a change in morphology (*i.e*., during the activation, there is an initial swelling of the spore that occurs before tube germination and can be seen one day after flushing CO_2_); and 2) a change in the contrast of the spore: a) the unactivated spores appear as bright, highly refractive bodies, similar to *Nosema* spp. spores using the same microscopy techniques (Tauber, Collins, et al., 2019), and b) the difference in the brightness of activated and unactivated spores appears similar to *Ascosphaera aggregata* (Stephen, Vandenberg, Fichter, & Lahm, 1982).

### Statistics

Within the hemocytometer we counted unactivated and activated spores in two separate 1 mm × 1 mm counting chambers per tube. Each treatment had two to four replicates that were run in parallel per trial and the experiment was repeated four times, totaling 24 replicate counts per treatment. We summed the counts per treatment across all trials for both activated and unactivated spores, and initially ran a Chi-squared test followed by 2×2 Fisher’s exact tests (treatment versus activated/unactivated). Calculations were done online using the tools provided on www.socscistatistics.com (accessed 2019). Unactivated spore proportions (number of unactivated spores over total spores counted in one counting chamber (1 counting chamber count = 1 data point)) between treatments were then compared. Based on these outputs, we proceeded to create a mixed model in order to compensate for possible random effects and to determine statistical significance between treatments of unactivated spore proportions. To this end, we created a generalized linear mixed model with a beta family distribution using default parameters as follows. First, 1e-6 was subtracted from every proportion before modeling. The model and statistics were calculated in RGui (3.6.1, 32-bit PC) using these packages: glmmTMB, effects, emmeans, multcomp, car, lme4 and glht. For the model, the unactivated spore count proportion was the dependent variable; the treatment was a fixed effect; and the trial, fungal glycerol stock preparation and dsRNA-T7 RNA polymerase reaction mixture (produced *in vitro* at two different times) were random variables. The final model included every noted factor. We independently tested each possible factor on treatment (described in the Results) for which interactions were found to be not statistically significant. Lastly, a multiple pairwise comparison of the model was done by the Tukey test with a Bonferroni correction. The alpha level was set at .05 in all cases and when *P* was less than .05 we considered the outcome to be statistically significant (*i.e*., we rejected that there was no difference/association between *As. apis*-targeting dsRNA and spore activation). Graphs were made online using the Plotly graphing platform available on chart-studio.plot.ly/ (accessed 2019).

## Results

We counted activated and unactivated *As. apis* spores that were incubated in the presence of Ras family or beta-glucan targeting dsRNA, unrelated dsRNA (control) or a mock control (blank water). The null hypothesis was that there was no association/difference between the first step in *As. apis* spore germination (*i.e*., activation) and the presence of dsRNA. The 5×2 Chi-squared test of the five treatments and two categories of activated or unactivated spore counts revealed a significant difference between one or more of the two treatments (χ^2^ = 6214.8899, df = 4, *P* < 0.00001), likely either due to lack of CO_2_ as expected for the negative control or from one of the dsRNA treatments (alternative hypothesis). 2×2 Fisher’s exact tests were then used where each treatment was compared to the positive control. Here, we observed statistically significant associations between CO_2_ (*i.e*., with the negative control) and the active state of the spore (*P* < 0.00001), as well as between the active state of the spore and each *As. apis*-targeting dsRNA (each, *P* < 0.00001). This was not observed, however, for the unrelated dsRNA control (*P* = .6740). Additionally, the average and median of unactivated spore proportions in the *As. apis* dsRNA Ras gene treatment (average =0.3031 and median =0.2964) and *As. apis* dsRNA beta-glucan gene treatment (=0.4026 and =0.3325) were higher than the positive control (=0.2572 and =0.2621) and also the unrelated dsRNA control (=0.2621 and =0.2545); all were lower than the negative control (=0.9893 and =0.9921).

Given these results, we chose to model and test the means of the groups because it appeared likely that the presence of *As. apis*-targeting dsRNA was responsible for the difference in proportions. We initially tested the model by checking for experimental effects on the treatments: the date that the dsRNA reaction mixtures were produced indicated no statistically significant interaction with the treatments (χ^2^ = 2.65, df = 4, *P* = .6188); the independent date that the trial was performed was not significant although a possible interaction with the treatment was observed (χ^2^ = 18.82, df = 12, *P* = .0930); and the fungal stock used had no statistically significant interaction with the treatments (χ^2^ = 6.00, df = 8, *P* = .6481).

We observed a statistically significant average higher proportion of unactivated spores when *As. apis* beta-glucan dsRNA was co-incubated with the spores compared to the positive control (*P* < .0001; Figure 1). The model-derived estimated marginal means computed with 95% confidence intervals confirmed that these two treatments did not overlap (positive control: = -1.054 ± 0.123 SE; CI_95%_ [-1.298, -0.811]; and *As. apis* dsRNA beta-glucan gene treatment: = -0.407 ± 0.118 SE; CI_95%_ [-0.642, -0.173]). We also removed seven non-statistically significant (via the Grubbs’ test), higher-than-average unactivated *As. apis* proportion data points from the *As. apis* beta-glucan gene dsRNA treatment. After removing these seven data points from the *As. apis* beta-glucan gene dsRNA treatment and rerunning the exact same statistical analyses, the resulting model-, multiple comparison (Tukey)-, Bonferroni-derived result remained statistically significant (Positive control vs. modified *As. apis* beta-glucan gene dsRNA treatment: *P* = .0057; Figure 1). The negative control that did not receive CO_2_ showed almost no activated spores the next day.

**Figure 1.**
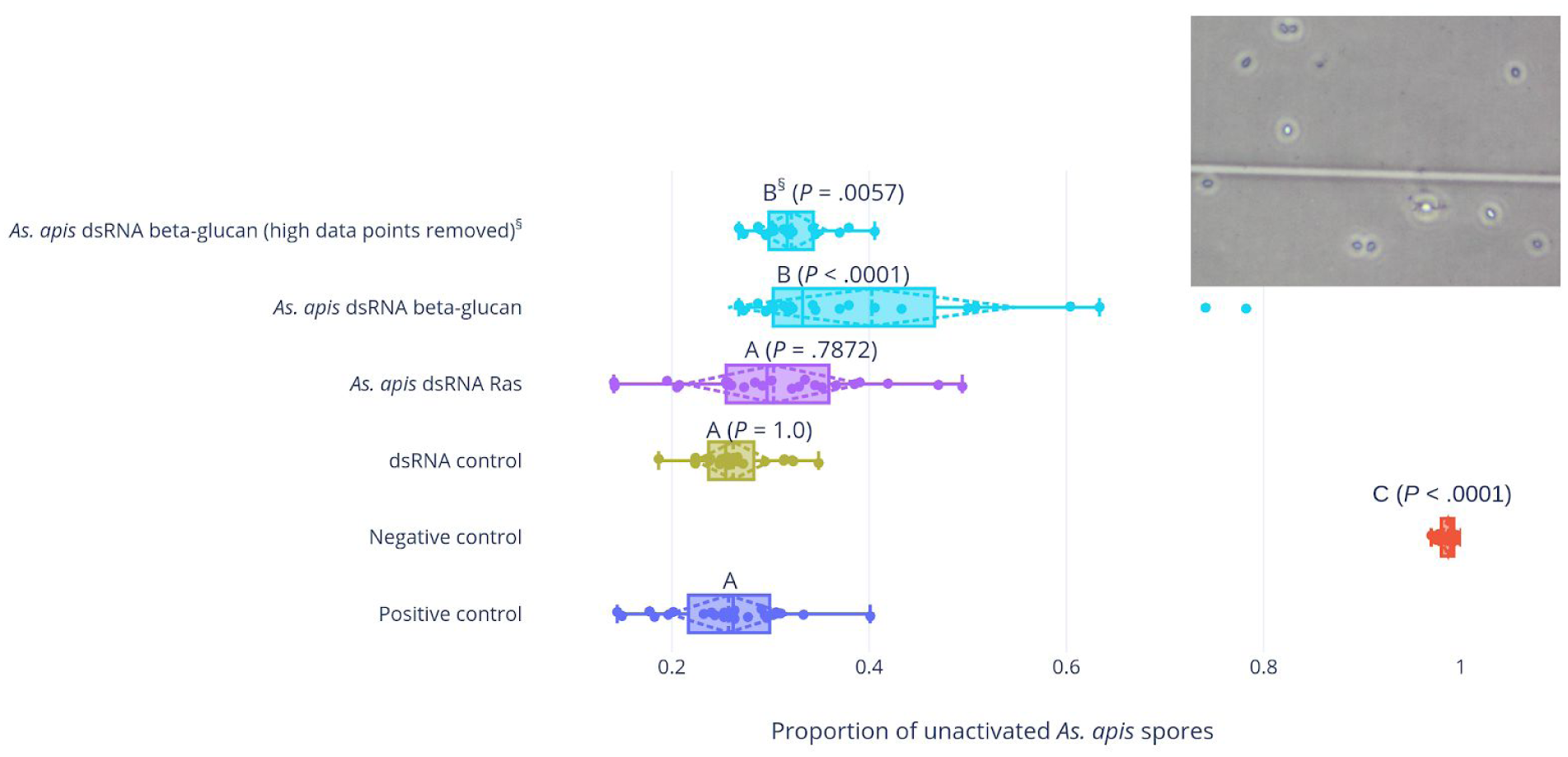
Box plots showing the median and quartiles of unactivated *As. apis* spore proportions. A scatter plot and dotted mean/standard deviation plot are overlaid. Above the box plot we show the resulting Bonferroni-corrected *P* value from the model-derived, Tukey multiple comparison test whereby each treatment was compared to the positive control. A different letter indicates a statistically significant difference of the treatment to the positive control. We observed a statistically significant higher proportion of unactivated spores when the *As. apis* spores were incubated with *As. apis* dsRNA targeting its beta-glucan gene. ^§^ Note: non-significant (via the Grubbs’ test) yet higher-than-average data points (seven data points in total) of unactivated spore proportions were removed from the *As. apis* beta-glucan gene dsRNA treatment dataset and the exact same model and statistical tests ran again. This was done to conservatively see if the conclusion remained statistically significant because these data points pulled the mean away from the median. For this ^§^ box plot, we show the resulting model-, multiple comparison (Tukey)-, Bonferroni-derived *P* value between this reduced dataset and the positive control while all of the other *P* values in the figure are from the former, full dataset statistical test. **Inset photograph:** Microscopy picture (400 × in phase contrast) of *As. apis* spores 24 hours after a CO_2_ flush. After 24 hours, in the treatments (except for the negative control) we observed a mixture of unactivated (bright, highly refractive bodies) and activated spores (dark bodies), the latter of which were swelling spores (a halo was observed around all the spores). For analysis, we used spore counts 24 hours after introducing CO_2_ and the dsRNA.

## Discussion

*Ascosphaera apis* is part of the honey bee pathosphere (Schwarz, Huang & Evans, 2015) and should be considered when attempting to combat the many pathogens and stressors undermining honey bee populations. *As. apis* is prevalent when screened by PCR-based surveys. For example, a comprehensive survey on honey bee products in Brazil noted that 73.17% of the tested samples were contaminated with *As. apis* (compared to 87.80% from *N. ceranae*) (Teixeira et al., 2018). Although the relative severity of *As. apis* is not considered to be as alarming as that of *N. ceranae*, DWV or *Paenibacillus larvae* (American foulbrood disease), its high prevalence, ability to cross-contaminate amongst bees (Pereira, Meeus, & Smagghe, 2019), deterrent effect for bee pollination (Yousefi & Fouks, 2019), and potential to inflict damage on both healthy as well as weakened colonies indicate a need to further develop *As. apis* control strategies.

Other honey bee RNAi experiments have also shown success in reducing pathogen loads (reviewed in (Brutscher & Flenniken, 2015)). RNAi strategies against viruses were successful, at least under controlled laboratory conditions. In two separate studies, dsRNA targeting DWV caused a reduction in DWV titers (Desai, Eu, Whyard, & Currie, 2012; Ryabov et al., 2014). Similarly, one study showed a reduction in IAPV after feeding IAPV-targeting dsRNA to bees (Maori et al., 2009). Interestingly, researchers noted a reduction in IAPV levels from a large-scale study using 160 hives and bulk feeding of RNAi molecules targeting IAPV (Hunter et al., 2010). For *N. ceranae*, knocking down the transcripts of polar tube protein 3 and incubating the bees with *Nosema* for ten days led to a reduction in counted spores (Rodríguez-García et al., 2018). This was also the case when targeting *N. ceranae Dicer* as well as ADP/ATP transporters (the latter, as measured by 16s rRNA transcripts) (Huang et al., 2019; Paldi et al., 2010). A recent study used dsRNA to target the gut parasite trypanosomatid, *Crithidia mellificae*, *ex vivo* (*i.e*., tested outside of a bee host) (de Sousa Pereira et al., 2019). The researchers selected a conserved, likely essential gene for the microorganism, produced dsRNA targeting this gene and introduced the dsRNA via three methods (electroporation into the cells, transfection by a reagent or simply adding the dsRNA to the medium containing the cells). They observed that there was a marginal reduction in growth rate when the dsRNA was simply added to the medium and this reduction was dependent on cell density. Our methods were similar to the *C. mellificae*-dsRNA experiment in that we simply added the dsRNA to the medium and then counted spores. The uptake of external RNA species during fungal growth was observed in *P. ruminantium* (activated spores), *A. fumigatus* (during outgrowth from activated spores), *A. nidulans* (activated and germinating spores), *Sclerotinia sclerotiorum* (mycelia outgrowth; (McLoughlin et al., 2018)) and *Botrytis cinerea* (germinating spores; (Wang et al., 2016)). Based on these experiments with fungi, the use of transfection reagents or a special delivery system appears unnecessary and that fungi will uptake various RNA species (single-stranded or double-stranded) of any sequence. Based on these studies and our own, we speculated that upon activation but before swelling, *As. apis* already began to uptake nutrients as well as exogenous RNA species. Additionally, these noted experiments tested whether the microorganism will selectively self-inflict damage, thus reducing the transcripts of the gene of interest. This was similar to our experiment where it appeared that the fungus negatively acted on itself causing the spore to not proceed to swell even after the introduction of CO_2_. We tested dsRNA targets against annotated Ras family protein and beta-glucan genes as these genes are putatively essential and lethal when knocked out, at least in other fungi. As noted in *Pecoramyces ruminantium*, uptake of exogenous dsRNA in activated spores can happen very quickly (measured in the study at 15 minutes from the onset of activation with potential that uptake happened sooner).

Moving forward, *in vivo* host tests require great care in order to reduce or eliminate any possible side effects of the dsRNA. On the one hand, ambiguous dsRNA sequences can already induce an RNAi antiviral defense response in the bee (Flenniken & Andino, 2013). On the other hand, researchers must carefully design their dsRNA sequences to mitigate off-target effects. For example, even the use of *GFP* sequences, which have limited similarity to the *A. mellifera* genome, can result in systemic unintended changes in gene expression in the host (Jarosch & Moritz, 2012; Nunes et al., 2013). Moreover, improved knockdown approaches such as delivery systems should not be discarded as they may further increase the efficiency of RNAi inhibition, and possible synergy with the bee host RNAi defense system (*i.e*., tested *in vivo*) may also improve the pathogen control outcomes. Although use of dsRNA *ex vivo* and *in vivo* has been successfully used to control DWV, IAPV, *N. ceranae*, *C. mellificae* and now *As. apis*, its success on a large scale, namely in a commercial setting, remains to be determined. Additionally, there is always an evolutionary arms race between combating the pathogen and escape of the pathogen from any selective pressure. For example, DWV appears to infect or replicate as a recombinant pool of genotypes, thus the current RNAi methods need to keep pace with the varying sequences of DWV (namely, researchers must be concerned that the chosen dsRNA sequence will lose efficacy as the complementary viral sequence changes, as already observed in bees (Ryabov et al., 2019)). Therefore, a combination of intelligently-designed, diverse and synergistic honey bee disease strategies from apicultural management practices to scientific advancements, will eventually and undoubtedly lead to improved honey bee-dependent agriculture. This will also improve our understanding of insect host immunity and virulence factors. Overall, we rejected our null hypothesis and concluded that the presence of *As. apis*-targeting dsRNA can negatively impact spore activation.

## Conclusion

*Ascosphaera apis* is a honey bee-associated fungal pathogen and as of yet there has not been an RNAi control strategy developed against this fungus. We found that if we incubated *As. apis* spores with *As. apis*-targeting dsRNA, then we observed a higher proportion of unactivated spores, suggesting that the presence of dsRNA inhibited, at least partially, the swelling/initial germination stage of the spores. Therefore, researchers may be able to exploit the RNAi mechanism of *As. apis*, and potentially in synergy with honey bee’s own RNAi machinery, to test fungal virulence factors and develop pathogen control strategies.

## Acknowledgements

J.T. was supported by an appointment to the Agricultural Research Service (ARS) Research Participation Program administered by the Oak Ridge Institute for Science and Education (ORISE) through an interagency agreement between the U.S. Department of Energy (DOE) and the U.S. Department of Agriculture (USDA). ORISE is managed by ORAU under DOE contract number DE-SC0014664. J.T. was also supported by the ARS Research Associate Program (Class of 2018). USDA is an equal opportunity provider and employer. The authors declare that they have no competing interests. All opinions expressed in this paper are the author’s and do not necessarily reflect the policies and views of USDA, ARS, DOE, or ORAU/ORISE. Mention of trade names or commercial products in this publication is solely for the purpose of providing specific information and does not imply recommendation or endorsement by the U.S. Department of Agriculture. We thank Baris Tursun and Anne Krause for supplying the bacterial strain and plasmid; Dawn Lopez for coordinating various aspects of this work; Kerstin Klutzny for supplying *S. lacrymans* from the BAM reference organism collection; Louela A. Castrillo and others at the USDA-ARS for organizing and shipping the fungal strains; Thorben Sieksmeyer for helpful discussions regarding the project; and Hannes Meissner for preparing what was needed for the experiments. J.T., D.M., R.E. and J.E. designed the project; J.T. did the experiments and analyses; J.T. wrote the manuscript with contributions from others; D.M. and J.E. funded/contributed reagents for this project; J.T., D.M., R.E. and J.E. approved the submitted manuscript.

## Supplement

**Sequences**:

>Ras gene (Sequenced cds (trimmed plasmid), primer bolded)

**CCTGCAATGGTTTGTGGACG**ACCCTTCAGAGTGTTGCCAGGGGTAATGCTGTGCG TGACAAGACCCAGGAAACTCGCTCAGAATTGCAGGAACAGGAACCATCTATCATCAGTCTGCAAAGCTCTATTAGGGGCGCTCTGCTACGTGACTGGGTGGATGATACACTCAGCGCATTGGACGAGAACCAGGACTCTATACTGCAATTCCAAGCTATCGCAAGAGGGCGACTGTGTCGACAGAAGATAGGCGAAGACCACCTGGACCTTCTTGCGGAGGAAGAGCAGATCACCAAATTCCAGGCGTTGGGACGAGGCGCTTTGCAGCGCGTTATCTTGGTGGATCTCTTTGAACAATTGGATGCATGTCTCCCCAAGATCGAAGAGCTA CAGTCTCTTGCTCGTG**GCAGGAACCTCAGGCTGAAT**

>beta-glucan gene (Sequenced cds (trimmed plasmid), primer bolded, reverse complement)

**TTTACCGTTGTTTGCGCCAG**AGACGTTGATTCCCAATCCACTACCCGCACTAGCCC TTTTTGCTGCAGCAGCAGCCTGAGCCTGCTGCTTGAAGGGCTTCAATTTCAACTCC\ CCGGAATCCTCATCGACGGAGACATGAGGCGATAAGAAATCGTCGGCGAGCATAG CGAGGAAGGAAGCCCATGATCGTGCAACGACGTATTTACAATCGAAGTCTCGTCC GAATATGATTACTTGGCCCCATTTCCCAGCTGGACCGGGCGCGAGATCAACGGCG ATATTGTTACCGCCCCAGTCCCGCGCAAGGGGGATCCATGACGGGTGAGCGTATG CCTTCTGCACCGCGCGCGGAGGGTGAGAGTCTTGTTTATCGAGTAACTGCTGTCT CCAGAGGGAATTGCCCTGGGACTGTAGCGTGGGCGTTGAGGCCCCCGA**GGAAGAAGTACCCACGCCAA**

## References

Baldwin, T., Islamovic, E., Klos, K., Schwartz, P., Gillespie, J., Hunter, S., & Bregitzer, P. (2018). Silencing efficiency of dsRNA fragments targeting Fusarium graminearum TRI6 and patterns of small interfering RNA associated with reduced virulence and mycotoxin production. PloS One, 13(8), e0202798. https://doi.org/10.1371/journal.pone.0202798

Brutscher, L. M., Daughenbaugh, K. F., & Flenniken, M. L. (2017). Virus and dsRNA-triggered transcriptional responses reveal key components of honey bee antiviral defense. Scientific Reports, 7(1), 6448. https://doi.org/10.1038/s41598-017-06623-z

Brutscher, L. M., & Flenniken, M. L. (2015). RNAi and Antiviral Defense in the Honey Bee. Journal of Immunology Research, 2015, 941897. https://doi.org/10.1155/2015/941897

Calkins, S. S., Elledge, N. C., Mueller, K. E., Marek, S. M., Couger, M. B., Elshahed, M. S., & Youssef, N. H. (2018). Development of an RNA interference (RNAi) gene knockdown protocol in the anaerobic gut fungus Pecoramyces ruminantium strain C1A. PeerJ, 6, e4276. https://doi.org/10.7717/peerj.4276

Cornman, R. S., Bennett, A. K., Murray, K. D., Evans, J. D., Elsik, C. G., & Aronstein, K. (2012). Transcriptome analysis of the honey bee fungal pathogen, Ascosphaera apis: implications for host pathogenesis. BMC Genomics, 13, 285. https://doi.org/10.1186/1471-2164-13-285

Dang, Y., Yang, Q., Xue, Z., & Liu, Y. (2011). RNA interference in fungi: pathways, functions, and applications. Eukaryotic Cell, 10(9), 1148–1155. https://doi.org/10.1128/EC.05109-11

de Sousa Pereira, K., Piot, N., Smagghe, G., & Meeus, I. (2019). Double-stranded RNA reduces growth rates of the gut parasite Crithidia mellificae. Parasitology Research, 118(2), 715–721. https://doi.org/10.1007/s00436-018-6176-0

Desai, S. D., Eu, Y.-J., Whyard, S., & Currie, R. W. (2012). Reduction in deformed wing virus infection in larval and adult honey bees (Apis mellifera L.) by double-stranded RNA ingestion. Insect Molecular Biology, 21(4), 446–455. https://doi.org/10.1111/j.1365-2583.2012.01150.x

Döke, M. A., Frazier, M., & Grozinger, C. M. (2015). Overwintering honey bees: biology and management. Current Opinion in Insect Science, 10, 185–193. https://doi.org/10.1016/j.cois.2015.05.014

Flenniken, M. L., & Andino, R. (2013). Non-specific dsRNA-mediated antiviral response in the honey bee. PloS One, 8(10), e77263. https://doi.org/10.1371/journal.pone.0077263

Goettel, M., Duke, G., Schaalje, G., & Richards, K. (1992). Effects of selected fungicides on in vitro spore germination and vegetative growth of Ascosphaera aggregata. Apidologie, 23(4), 299–309.

Heath, L. A. F., & Gaze, B. M. (1987). Carbon Dioxide Activation of Spores of the Chalkbrood Fungus Ascosphaera Apis. Journal of Apicultural Research, 26(4), 243–246. https://doi.org/10.1080/00218839.1987.11100768

Hu, Y., Stenlid, J., Elfstrand, M., & Olson, Å. (2013). Evolution of RNA interference proteins dicer and argonaute in Basidiomycota. Mycologia, 105(6), 1489–1498. Retrieved from JSTOR.

Huang, Q., Li, W., Chen, Y., Retschnig-Tanner, G., Yanez, O., Neumann, P., & Evans, J. D. (2019). Dicer regulates Nosema ceranae proliferation in honeybees. Insect Molecular Biology, 28(1), 74–85. https://doi.org/10.1111/imb.12534

Hunter, W., Ellis, J., Vanengelsdorp, D., Hayes, J., Westervelt, D., Glick, E., … Paldi, N. (2010). Large-scale field application of RNAi technology reducing Israeli acute paralysis virus disease in honey bees (Apis mellifera, Hymenoptera: Apidae). PLoS Pathogens, 6(12), e1001160. https://doi.org/10.1371/journal.ppat.1001160

Jarosch, A., & Moritz, R. F. A. (2012). RNA interference in honeybees: off-target effects caused by dsRNA. Apidologie, 43(2), 128–138. https://doi.org/10.1007/s13592-011-0092-y

Jasrapuria, S., Specht, C. A., Kramer, K. J., Beeman, R. W., & Muthukrishnan, S. (2012). Gene families of cuticular proteins analogous to peritrophins (CPAPs) in Tribolium castaneum have diverse functions. PloS One, 7(11), e49844. https://doi.org/10.1371/journal.pone.0049844

Jensen, A. B., Aronstein, K., Flores, J. M., Vojvodic, S., Palacio, M. A., & Spivak, M. (2013). Standard methods for fungal brood disease research. Journal of Apicultural Research, 52(1), 1–20. https://doi.org/10.3896/IBRA.1.52.1.13

Jöchl, C., Loh, E., Ploner, A., Haas, H., & Hüttenhofer, A. (2009). Development-dependent scavenging of nucleic acids in the filamentous fungus Aspergillus fumigatus. RNA Biology, 6(2), 179–186. https://doi.org/10.4161/rna.6.2.7717

Kalleda, N., Naorem, A., & Manchikatla, R. V. (2013). Targeting fungal genes by diced siRNAs: a rapid tool to decipher gene function in Aspergillus nidulans. PloS One, 8(10), e75443. https://doi.org/10.1371/journal.pone.0075443

Maori, E., Paldi, N., Shafir, S., Kalev, H., Tsur, E., Glick, E., & Sela, I. (2009). IAPV, a bee-affecting virus associated with Colony Collapse Disorder can be silenced by dsRNA ingestion. Insect Molecular Biology, 18(1), 55–60. https://doi.org/10.1111/j.1365-2583.2009.00847.x

McLoughlin, A. G., Wytinck, N., Walker, P. L., Girard, I. J., Rashid, K. Y., de Kievit, T., … Belmonte, M. F. (2018). Identification and application of exogenous dsRNA confers plant protection against Sclerotinia sclerotiorum and Botrytis cinerea. Scientific Reports, 8(1), 7320. https://doi.org/10.1038/s41598-018-25434-4

Nunes, F. M. F., Aleixo, A. C., Barchuk, A. R., Bomtorin, A. D., Grozinger, C. M., & Simões, Z. L. P. (2013). Non-Target Effects of Green Fluorescent Protein (GFP)-Derived Double-Stranded RNA (dsRNA-GFP) Used in Honey Bee RNA Interference (RNAi) Assays. Insects, 4(1), 90–103. https://doi.org/10.3390/insects4010090

Paldi, N., Glick, E., Oliva, M., Zilberberg, Y., Aubin, L., Pettis, J., … Evans, J. D. (2010). Effective gene silencing in a microsporidian parasite associated with honeybee (Apis mellifera) colony declines. Applied and Environmental Microbiology, 76(17), 5960–5964. https://doi.org/10.1128/AEM.01067-10

Pereira, K. de S., Meeus, I., & Smagghe, G. (2019). Honey bee-collected pollen is a potential source of Ascosphaera apis infection in managed bumble bees. Scientific Reports, 9(1), 4241. https://doi.org/10.1038/s41598-019-40804-2

Rodríguez-García, C., Evans, J. D., Li, W., Branchiccela, B., Li, J. H., Heerman, M. C., … Chen, Y. P. (2018). Nosemosis control in European honey bees, Apis mellifera, by silencing the gene encoding Nosema ceranae polar tube protein 3. Journal of Experimental Biology, 221(19), jeb184606. https://doi.org/10.1242/jeb.184606

Ryabov, E. V., Childers, A. K., Lopez, D., Grubbs, K., Posada-Florez, F., Weaver, D., … Evans, J. D. (2019). Dynamic evolution in the key honey bee pathogen deformed wing virus: Novel insights into virulence and competition using reverse genetics. PLOS Biology, 17(10), e3000502. https://doi.org/10.1371/journal.pbio.3000502

Ryabov, E. V., Wood, G. R., Fannon, J. M., Moore, J. D., Bull, J. C., Chandler, D., … Evans, D. J. (2014). A Virulent Strain of Deformed Wing Virus (DWV) of Honeybees (Apis mellifera) Prevails after Varroa destructor-Mediated, or In Vitro, Transmission. PLoS Pathogens, 10(6). https://doi.org/10.1371/journal.ppat.1004230

Schmitt-Engel, C., Schultheis, D., Schwirz, J., Ströhlein, N., Troelenberg, N., Majumdar, U., … Bucher, G. (2015). The iBeetle large-scale RNAi screen reveals gene functions for insect development and physiology. Nature Communications, 6, 7822. https://doi.org/10.1038/ncomms8822

Shang, Y., Xiao, G., Zheng, P., Cen, K., Zhan, S., & Wang, C. (2016). Divergent and Convergent Evolution of Fungal Pathogenicity. Genome Biology and Evolution, 8(5), 1374–1387. https://doi.org/10.1093/gbe/evw082

Solis, C. F., Santi-Rocca, J., Perdomo, D., Weber, C., & Guillén, N. (2009). Use of bacterially expressed dsRNA to downregulate Entamoeba histolytica gene expression. PloS One, 4(12), e8424. https://doi.org/10.1371/journal.pone.0008424

Stephen, W. P., Vandenberg, J. D., Fichter, B. L., & Lahm, G. (1982). Inhibition of chalkbrood spore germination in vitro (Ascosphaera aggregata: Ascosphaerales).

Tauber, J. P., Collins, W. R., Schwarz, R. S., Chen, Y., Grubbs, K., Huang, Q., … Evans, J. D. (2019). Natural Product Medicines for Honey Bees: Perspective and Protocols. Insects, 10(10), 356. https://doi.org/10.3390/insects10100356

Tauber, J. P., Nguyen, V., Lopez, D., & Evans, J. D. (2019). Effects of a Resident Yeast from the Honeybee Gut on Immunity, Microbiota, and Nosema Disease. Insects, 10(9). https://doi.org/10.3390/insects10090296

Teixeira, É. W., Guimarães-Cestaro, L., Alves, M. L. T. M. F., Message, D., Martins, M. F., Luz, C. F. P. da, & Serrão, J. E. (2018). Spores of Paenibacillus larvae, Ascosphaera apis, Nosema ceranae and Nosema apis in bee products supervised by the Brazilian Federal Inspection Service. Revista Brasileira de Entomologia, 62(3), 188–194. https://doi.org/10.1016/j.rbe.2018.04.001

Wang, M., Weiberg, A., Lin, F.-M., Thomma, B. P. H. J., Huang, H.-D., & Jin, H. (2016). Bidirectional cross-kingdom RNAi and fungal uptake of external RNAs confer plant protection. Nature Plants, 2, 16151. https://doi.org/10.1038/nplants.2016.151

Yang, D., Xu, X., Zhao, H., Yang, S., Wang, X., Zhao, D., … Hou, C. (2018). Diverse Factors Affecting Efficiency of RNAi in Honey Bee Viruses. Frontiers in Genetics, 9, 384. https://doi.org/10.3389/fgene.2018.00384

Ye, J., Coulouris, G., Zaretskaya, I., Cutcutache, I., Rozen, S., & Madden, T. L. (2012). Primer-BLAST: a tool to design target-specific primers for polymerase chain reaction. BMC Bioinformatics, 13, 134. https://doi.org/10.1186/1471-2105-13-134

Yousefi, B., & Fouks, B. (2019). The presence of a larval honey bee parasite, Ascosphaera apis, on flowers reduces pollinator visitation to several plant species. Acta Oecologica, 96, 49–55. https://doi.org/10.1016/j.actao.2019.03.006

